## Supplementary Figures and legend for "How do bacterial extracellular Contractile Injection Systems bind target cells? A remarkable diversity of receptor binding domains": supp. figure legends.docx

**Supplementary Figure 1.** Computational pipeline for the identification and characterization of eCIS tail fiber domains. Schematic representation of the domain discovery workflow used to identify and characterize eCIS tail fiber proteins. The pipeline consists of four main stages:

1. Initial sequence collection from the eCIStem database (1,425 operons) and identification of 629 putative fiber genes through similarity to known Afp13/Pvc13 homologs.

2. Domain detection phase using all-against-all BLAST followed by clustering (40-20% identity thresholds) with CD-HIT and MMseqs2.

3. Multiple sequence alignments were generated using Clustal Omega and converted to Hidden Markov Models (HMMs) with HMMER.

4. Dataset expansion using N-terminal conserved domains in jackhmmer iterative searches against our comprehensive microbial genome database, increasing coverage to 3,445 fiber genes across 2,585 operons.

**Supplementary Figure 2**. Genomic organization and evolutionary patterns of multi-fiber eCIS operons

**(a)** Stacked bar chart showing the distribution of fiber gene counts per eCIS operon across the dataset (N=2,585 operons). Nearly one-quarter of operons (23.4%) encode multiple tail fibers, with a maximum of five fiber genes per locus.

**(b)** Representative genomic neighborhoods of eCIS operons containing 1-5 fiber genes. Fiber genes (deep red) consistently localize downstream of conserved baseplate components (AFP11\12 identified by Baseplate_J and GPW Pfams respectively and the tail_P2_I harboring genes).

**(c)** Example operon containing three structurally similar fiber genes with high sequence divergence (maximal 35% pairwise sequence identity). AlphaFold2 predictions reveal conserved eBAP2 with similarly looking fibers with C-terminal S74 domains.

**(d)** Operon encoding three architecturally distinct fibers: a common eBAP3 domain found on three fibers with diverged domain architectures.

**Supplementary Figure 3.** Structural basis of eBAPs fiber attachment to eCIS baseplate components

**(a-c)** Predicted trimeric structure of unexplored eBAPs with baseplate components: **a.** the AFP11-homolog (baseplate_J protein) (olive green) in complex with eBAP2 fiber N-terminal domains (pinkish).

**(b)**. eBAP3 domain in complex with Tail_P2_I harboring gene **c**. eBAP5 domain in complex with upstreen gene structurally similar to Tail_P2_I protein

**(d-f)** Zoomed view of the eBAPs baseplate interface. Key interactions include: eBAPs are predicted to interact with loops stemming from base plate proteins

**(g-i)** ipTM, pTM and Predicted Aligned Error (PAE) plots from AlphaFold3 multimer predictions.

**Supplementary Figure 4.** Evidence suggesting acquisition of domains in eCIS tail fibers

**(a)** Domain architecture of eBAP4 genes displays a mix of Ig-like domains ordered in different combinations of c-terminal chains. Domains detected on each gene by Pfam hmmscan displayed as domain architectures ordered from left to right (on amino acid sequence from N’ to C’ termini respectively). Gene ID displayed above each line.

**(b)** Taxonomic distribution of 64 Pfam domains showing phylogenetic incongruence across kingdoms. Stacked bars represent the relative share of Pfam group members from Bacteria (), Eukaryota-Metazoa (Dark red), Eukaryota-N\A (no detectable subgroup, purple), Eukaryota-Fungi (Olive green), Eukaryota-Viridiplantae (orange), Viruses (Navy blue) and Archaea (Yellow).These domains are with >90% non-bacterial representation suggest horizontal gene transfer events.

**(c)** C1q domains MSA from figure 3g.

**(d)** Adenovirus shaft homolog analysis. Top: Structural alignment demonstrating conservation of the triple β-spiral fold and key proline residues that maintain characteristic bend angles across viral and bacterial domains. Bottom: Phylogenetic tree showing bacterial eCIS fiber shaft domains clustering with viral sequences rather than forming a separate bacterial clade.

**Supplementary Figure 5.** Structural prediction pipeline and discovered domains network

**(a)** Schematic representation of the structural pipeline. We used a fasta database clustered by 70% similarity and coverage. Predicted trimeric structures of >1000 proteins. Used in two pathways. 1. Clustering of whole fibers. 2. Domain dissection via 3D models: secondary structures were extracted from PDB files we analyzed interactions between secondary structures by the rule of one closest residue. We clustered secondary structures in order to define domain boundaries. This resulted in 1,177 domain foldseek clusters.

**(b)** We generated a network of domain foldseek clusters with node size representing domain abundance. Coloring represents domain relative position on protein linear sequence. Edges are drawn if the clusters contain the same protein ids implying for shared architectures.

**Supplementary Figure 6.** Shoulder domains tend to get intertwined with adjacent structural features

Examples for eBAP3 **(a)** and eBAP4 **(b)** intertwining with downstream adjacent structural features. We used a rainbow diagram to demonstrate the fold’s back bone being included in the adjacent domain then going back and serving as beta-strands incorporated in the “shoulders” region typical to eBAP3-5. We observed that this trait, speculated to enhance structure stability, also enhances divergence in these protein families.

**Supplementary Figure 7.** Target recognition by screened eCIS tail fibers.

**(a)** Cell orientation of the fiberPb-modified PVC complex. Killing of A549, HeLa, and HEK293T cells by PVC complexes at 0.5 mg/mL concentration after 48h. Empty R7PVC was used as control.

**(b)** The possible fiber fragments in the predicted fiber proteins. Yellow parts represent the possible fiber fragments. Mr, Mycetohabitans rhizoxinica HKI 454; Am, Aquimarina sp. AU119. **(c)** Western blotting validation of engineered PVC assembly and protein loading. (D) The NSEM observation of engineered PVC particles. Scale bar, 100 nm. **(E-H)** Killing of THP-1, HEK293T, A549, and HeLa cells by engineered PVC complexes at 0.5 mg/mL concentration after 48h. Empty R7PVC was used as negative control.

**Supplementary Figure 8.** Verification of glycans’ effects on fiber-cell recognition.

**(a-c)** Glycans that cannot inhibit the fiberPb-PVC-cell recognition. Killing of THP-1 cells by 0.5 mg/mL PVC complexes pretreated with N-acetylglucosamine, D-Galactose or D-Fucose after 48h.

**(d-f)** D-Mannose cannot inhibit the cytotoxicity of R7PVC-TcsT. Killing of THP-1, HeLa, and HEK293T cells by 0.5 mg/mL R7PVC complexes pretreated with D-Mannose after 48h.
