## Supplementary figures and images for "How do bacterial extracellular Contractile Injection Systems bind target cells? A remarkable diversity of receptor binding domains"

### supp. fig 1.pdf

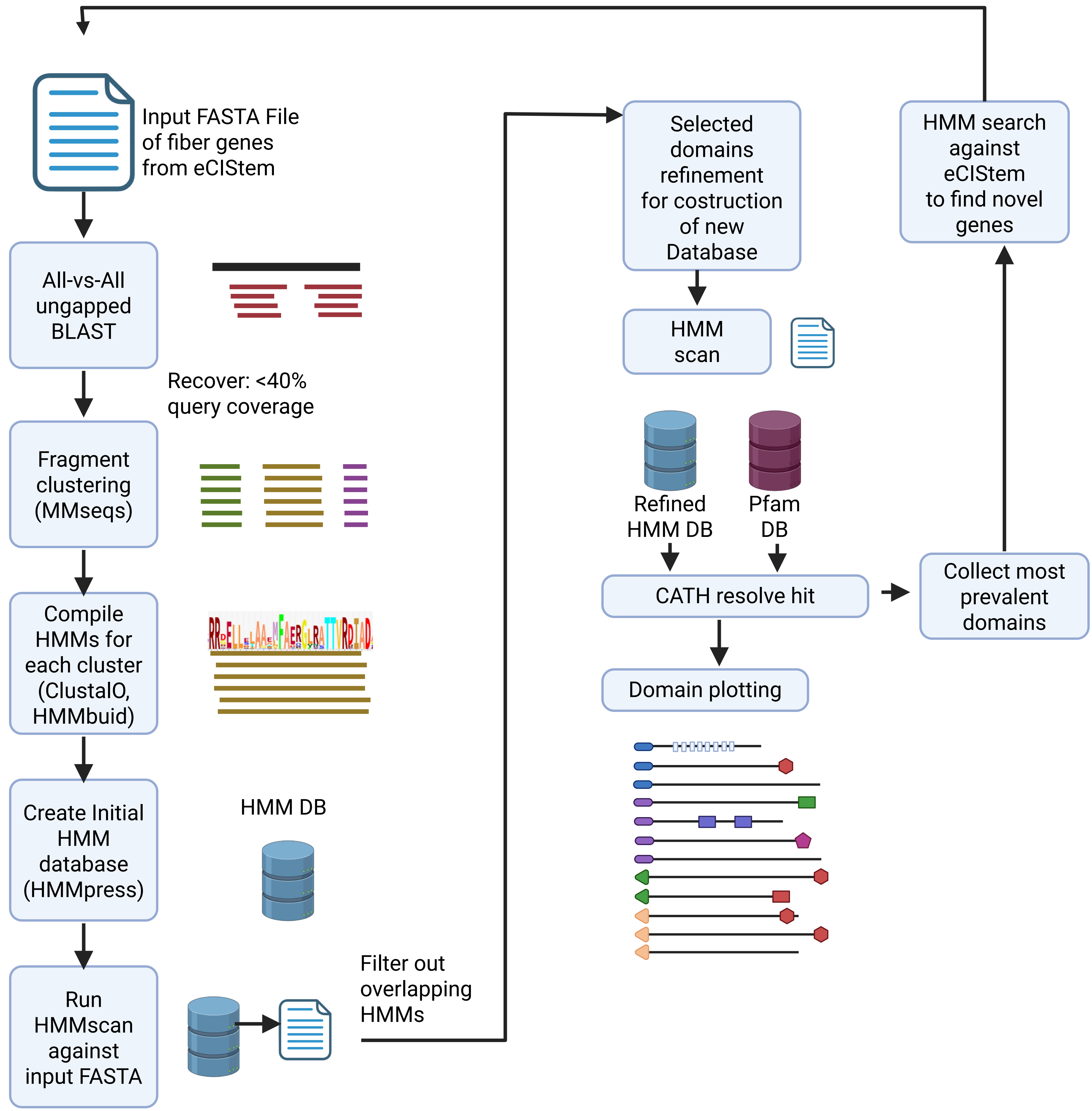

### supp. fig 2.pdf

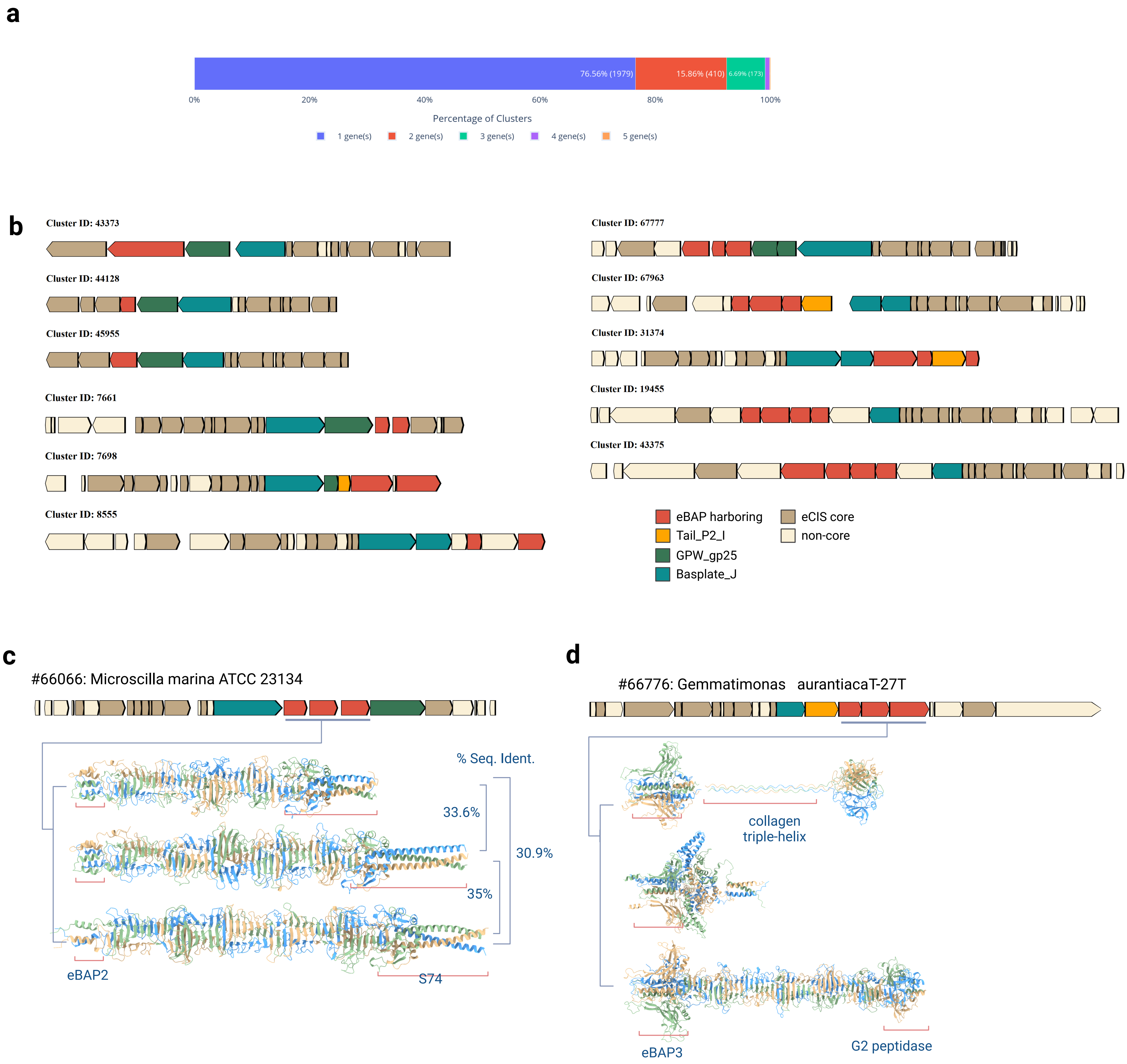

### supp. fig 3.pdf

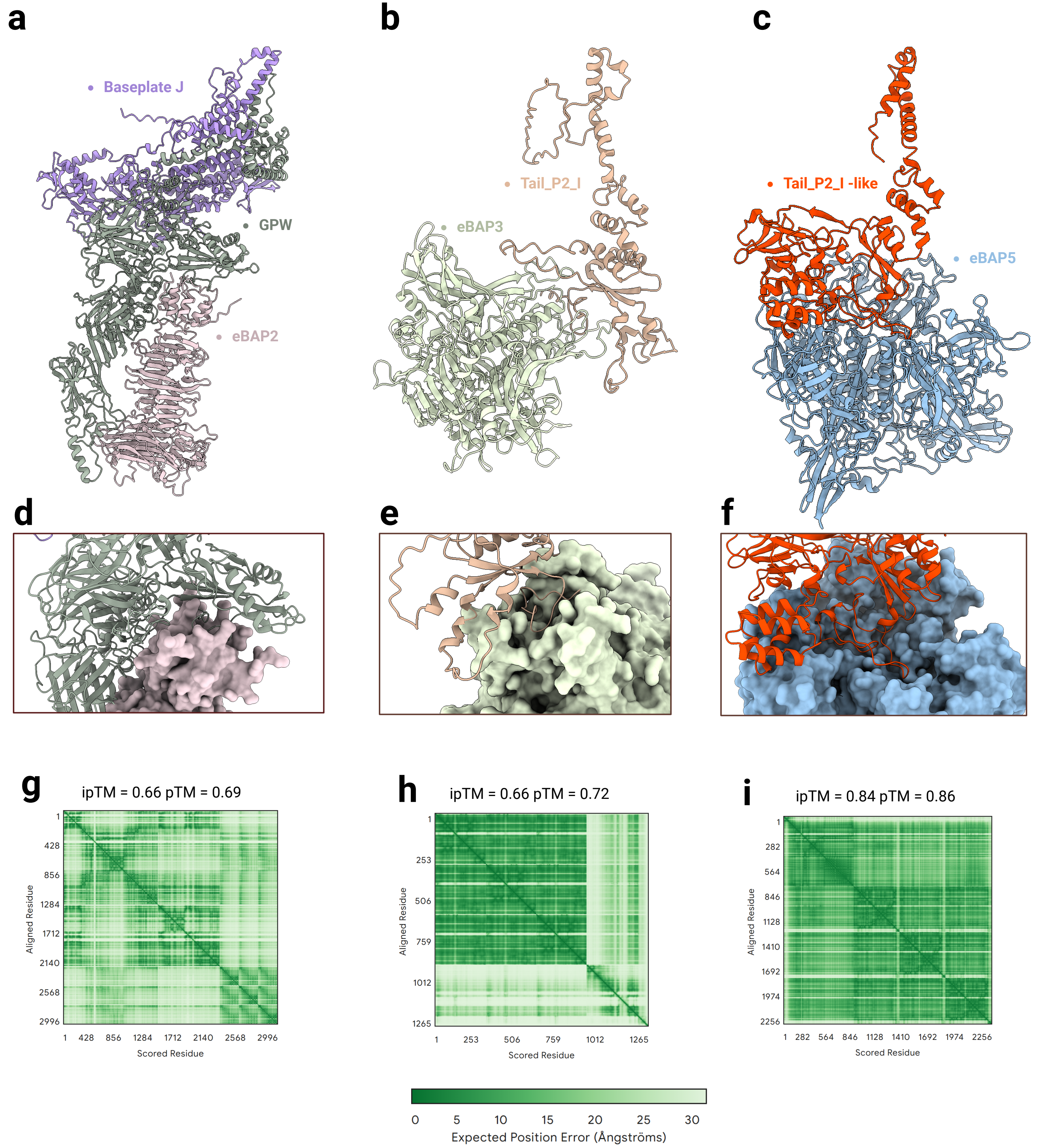

### supp. fig 4.pdf

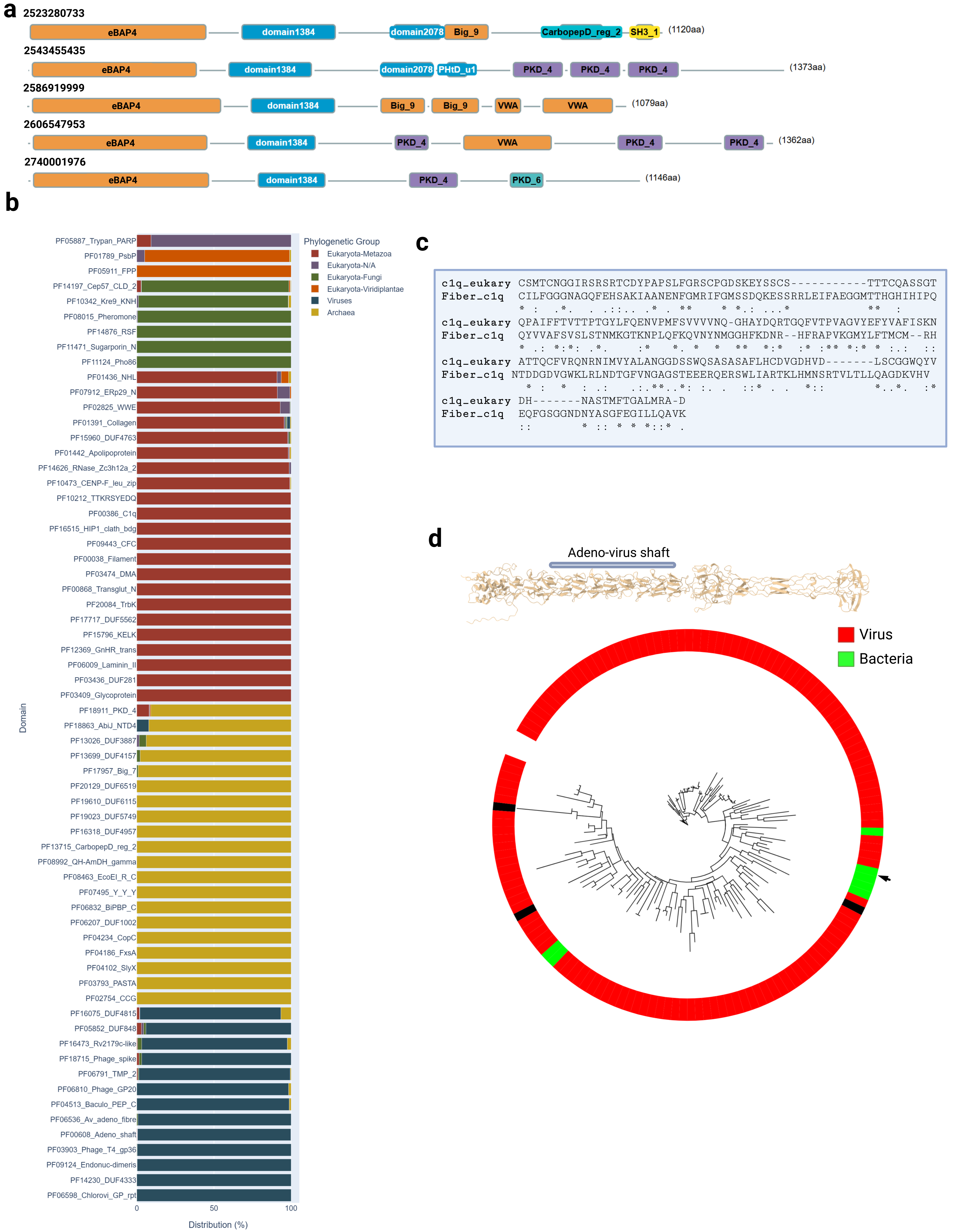

### supp. fig 5.pdf

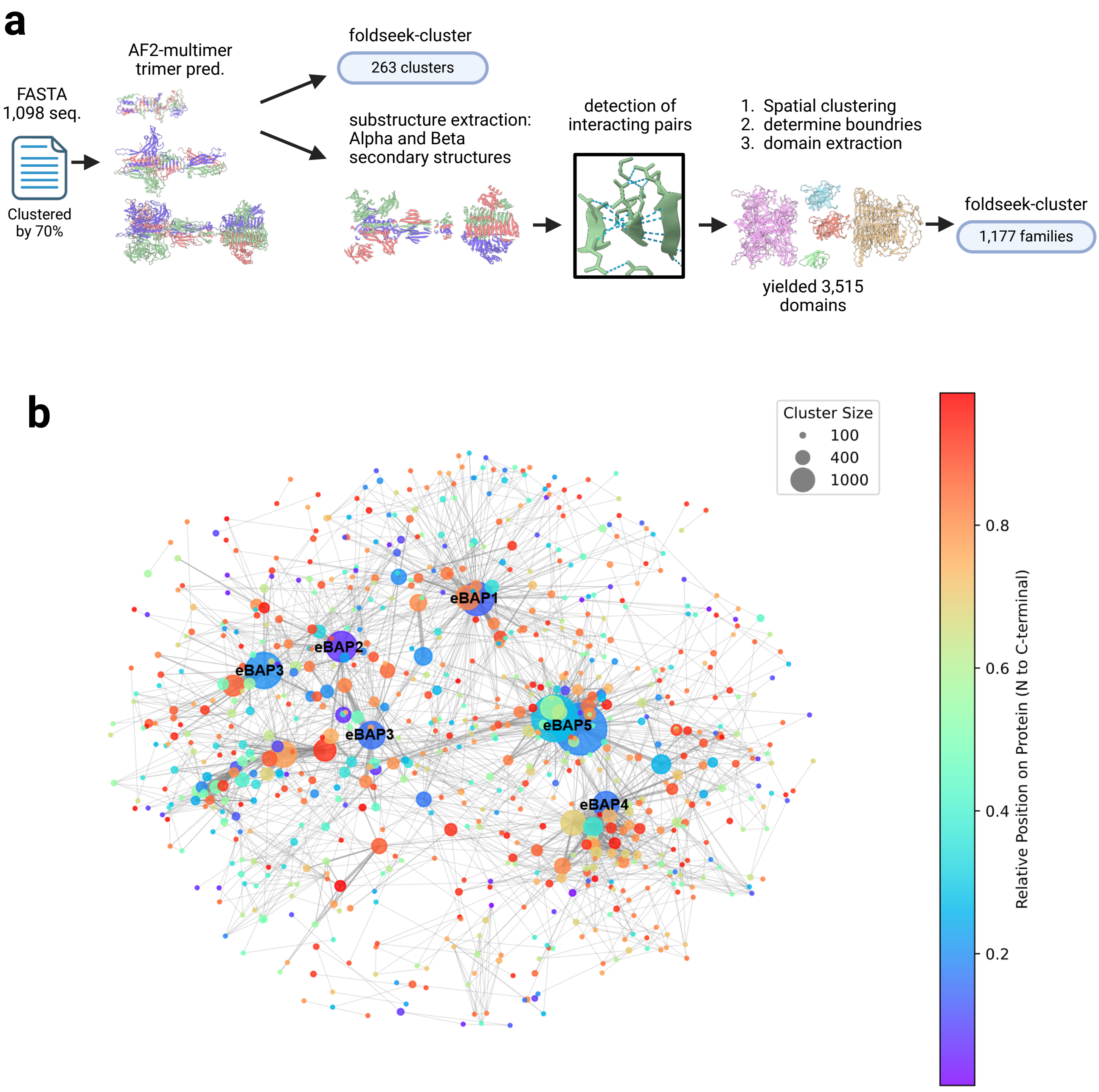

### supp. fig 6.pdf

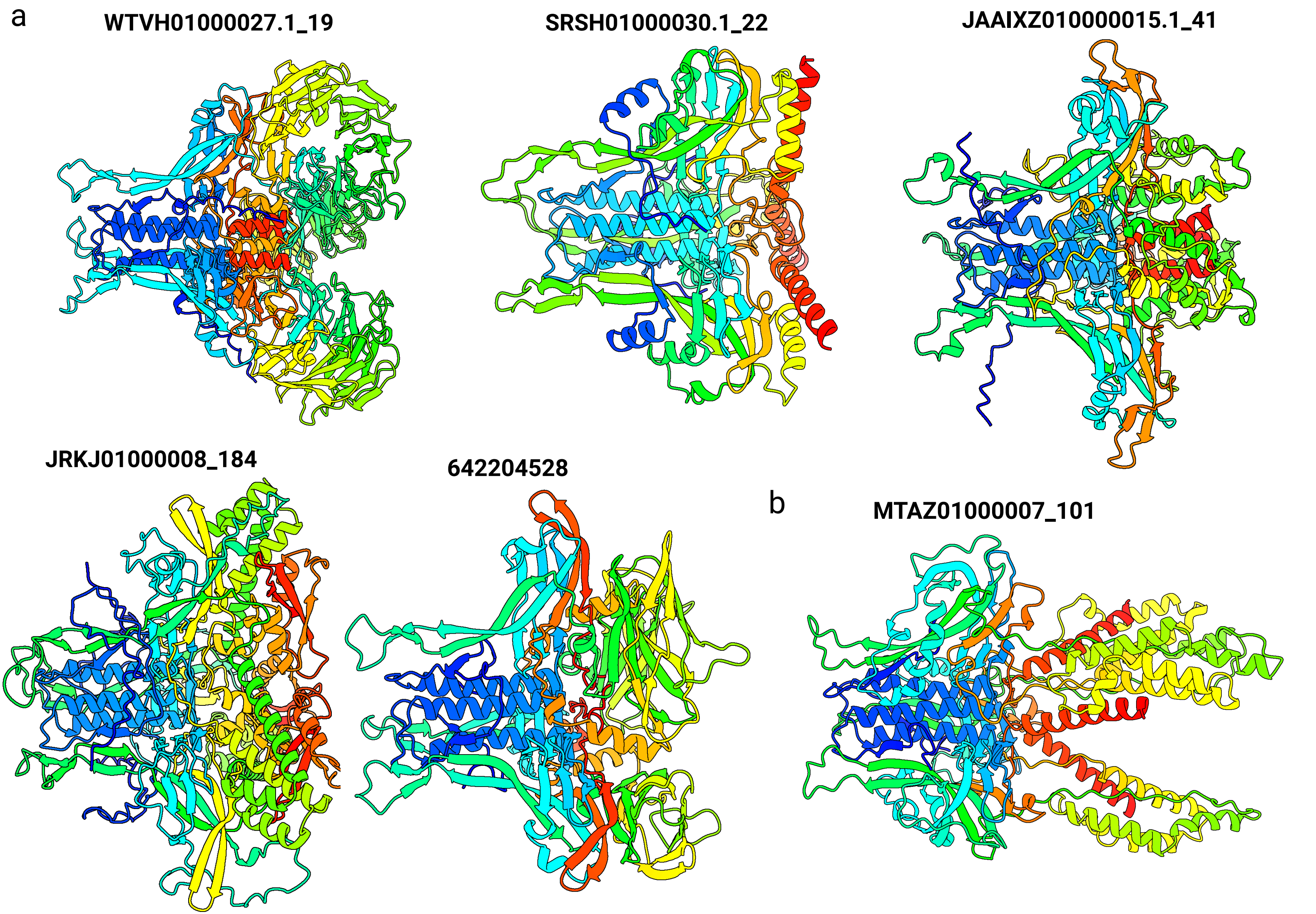

### supp. fig 7.pdf

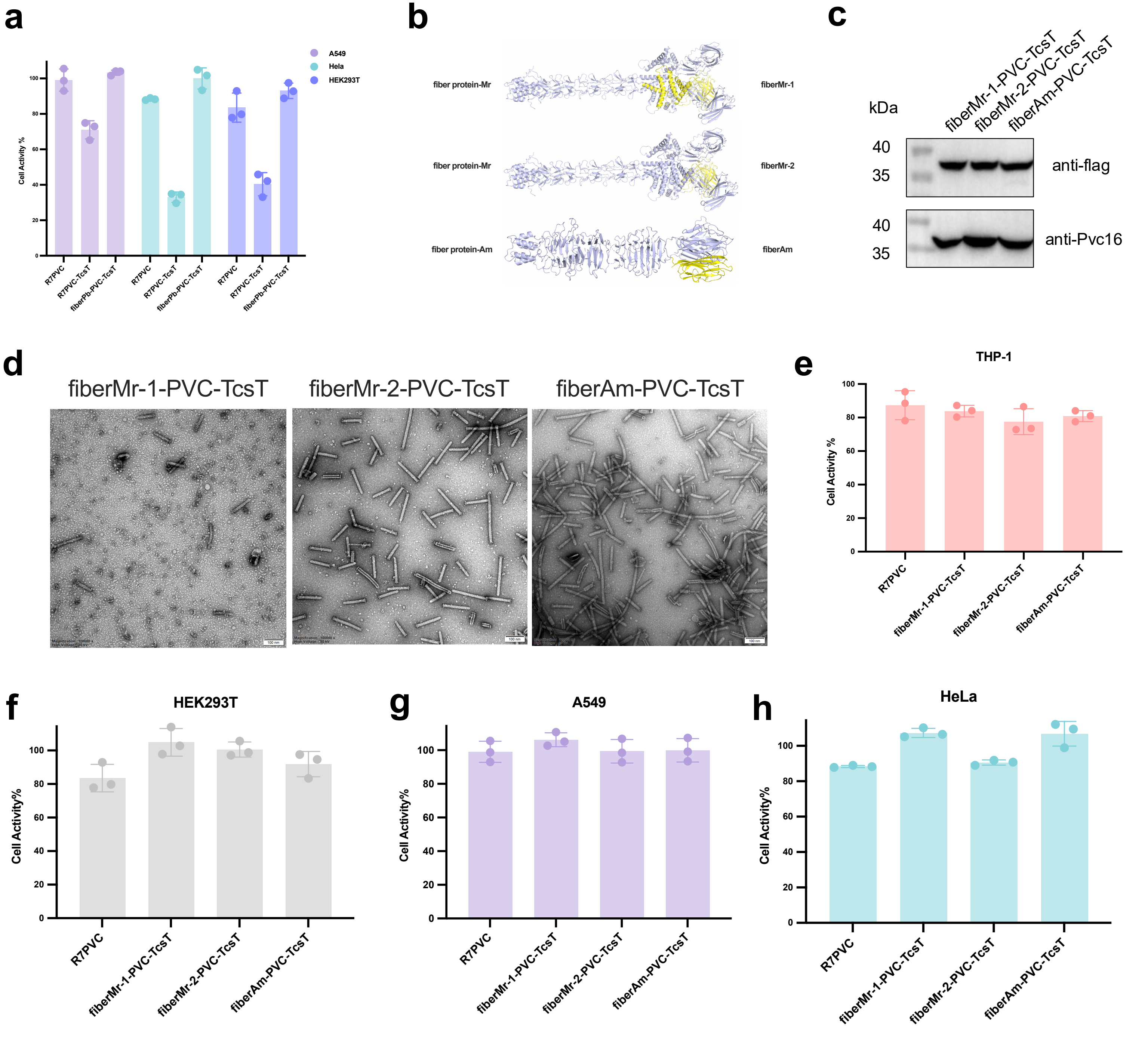

### supp. fig 8.pdf

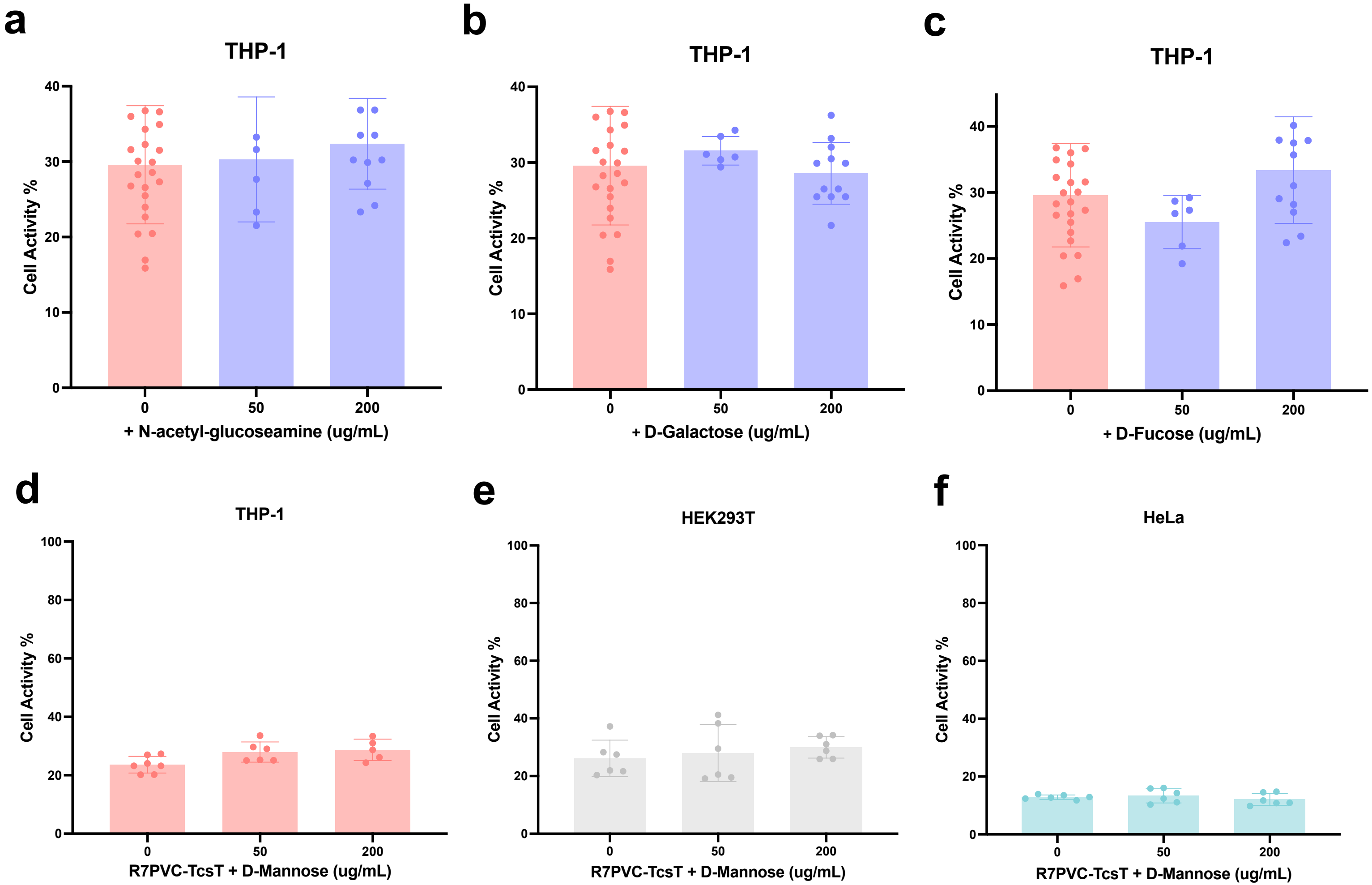
